# MicroRNA combinations function as synergistic network regulators of neuroblastoma differentiation

**DOI:** 10.64898/2026.03.04.709166

**Authors:** Seth A Lawson, Yiqiang Zhang, Adam Kosti, Matthew J Hart, Luiz O F Penalva, Alexander Pertsemlidis

## Abstract

Differentiation-based therapies represent a promising strategy for the treatment of neuroblastoma; however, single-agent approaches frequently yield incomplete and transient responses due to the robustness of underlying gene regulatory networks. MicroRNAs (miRNAs) are endogenous regulators of gene expression that modulate entire gene programs rather than individual molecular targets, making them attractive candidates for network-level therapeutic intervention. While individual miRNAs have been investigated as therapeutic agents, the potential for synergistic interactions between miRNAs remains largely unexplored.

Here, we developed a scalable high-content phenotypic screening platform to identify synergistic miRNA combinations that promote neuronal differentiation and growth arrest in neuroblastoma cells. Using SK-N-BE(2)-C cells and automated quantification of neurite outgrowth and confluence, we screened pairwise combinations of differentiation-associated miRNAs at submaximal doses. Candidate synergistic interactions were identified using the Highest Single Agent framework and subsequently validated by dose–response interaction modeling.

We identified a robust synergistic interaction between miR-124-3p and miR-363-3p that exceeded zero-interaction potency expectations by approximately 20.9% and increased maximal differentiation-associated phenotypic response by 73% relative to single-miRNA treatments. Target gene and pathway enrichment analyses revealed that miR-124-3p and miR-363-3p regulate largely distinct but functionally complementary target gene sets. These complementary targets converged on neuronal differentiation and cell cycle control pathways, providing a mechanistic basis for their cooperative activity.

Together, these findings establish miRNA combinations as programmable network regulators capable of inducing complex cellular phenotypes with greater efficacy than single agents. This work provides a conceptual and experimental framework for the rational discovery of synergistic miRNA therapeutics and suggests new avenues for differentiation-based treatment strategies in neuroblastoma and other diseases driven by dysregulated regulatory networks.

## 1. Introduction

Neuroblastoma is the most common extracranial solid tumor of childhood and remains a leading cause of cancer-related mortality in pediatric patients (Maris et al., 2007; Brodeur, 2003; Matthay et al., 2016). Although intensive multimodal therapy has improved outcomes for subsets of patients, long-term survival for high-risk disease remains limited and is accompanied by substantial treatment-related toxicity (Matthay et al., 1999; Matthay et al., 2009; Pinto et al., 2015). A defining biological feature of neuroblastoma is its capacity for neuronal differentiation, a property that has been therapeutically exploited through the use of retinoids as maintenance therapy following cytotoxic treatment (Reynolds et al., 2003; Matthay et al., 2009). However, differentiation-based strategies using single agents yield incomplete and often transient responses, reflecting the underlying complexity and redundancy of the gene regulatory networks that govern neuronal fate decisions (Chlapek et al., 2018; Stallings et al., 2011).

MicroRNAs (miRNAs) are endogenous small noncoding RNAs that regulate gene expression post-transcriptionally by targeting complementary sequences in messenger RNAs (He and Hannon, 2004; Bartel, 2009). Individual miRNAs typically modulate dozens to hundreds of transcripts, enabling them to act as higher-order regulators of cellular state rather than single-pathway effectors (Lewis et al., 2005; Sood et al., 2006). In neuroblastoma, multiple miRNAs have been implicated in differentiation, proliferation, and apoptosis, including miR-124, miR-34a, and miR-363 (Welch et al., 2007; Chen and Stallings, 2007; Buechner et al., 2011; Tivnan et al., 2014; Nolan et al., 2020). Despite this promise, therapeutic development of miRNAs has largely focused on single-miRNA interventions, which often produce modest phenotypic effects and exhibit limited durability (Nana-Sinkam and Croce, 2013; van Rooij and Kauppinen, 2014).

Combination therapy has long been recognized as a powerful strategy to overcome biological robustness by targeting multiple regulatory nodes simultaneously (Al-Lazikani et al., 2012). In the context of miRNAs, combinatorial regulation offers a unique opportunity to exploit the natural architecture of gene regulatory networks. Distinct miRNAs possess partially overlapping but non-identical target repertoires, suggesting that carefully selected miRNA pairs could cooperatively suppress complementary subsets of genes required to maintain malignant phenotypes (**Figure 1**) (Lewis et al., 2005; Agarwal et al., 2015). Such combinations may achieve phenotypic outcomes that exceed the sum of their individual effects, a property referred to as synergy (Bliss, 1939; Chou, 2006; Slinker, 1998).

**Figure 1.**
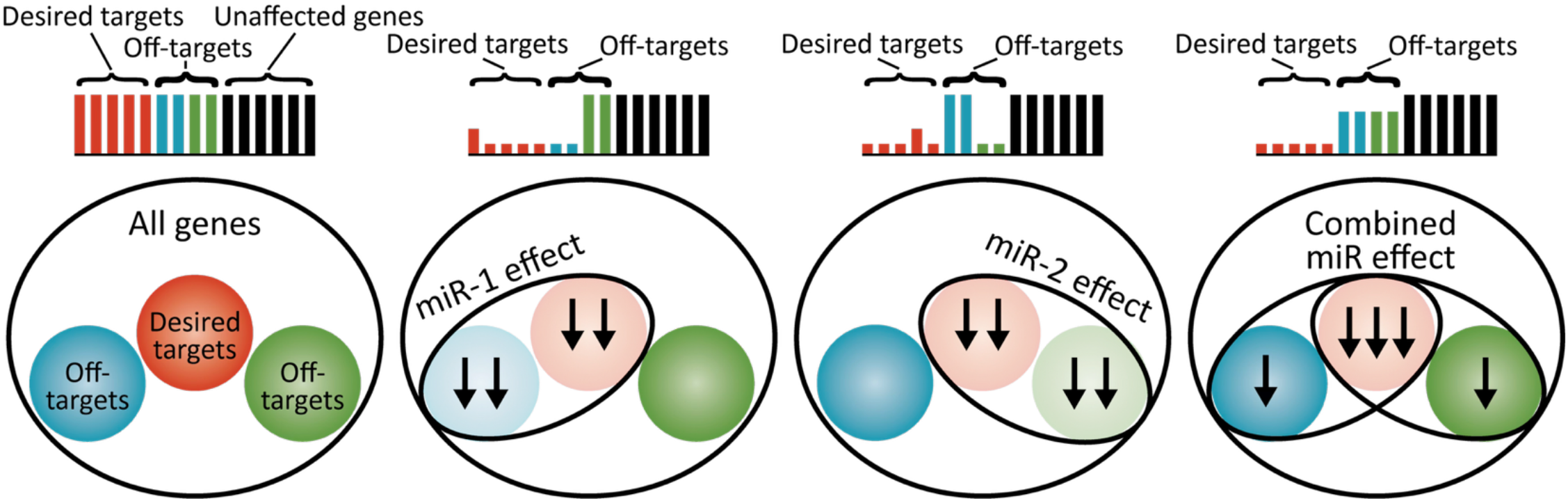
The overall target space includes desired targets (maroon) and (undesired) off-targets (blue and green). An individual miRNA down-regulates both desired targets and off-targets. A carefully selected combination of mimics can act synergistically to enhance the efficacy of NB differentiation while reducing the undesirable, or ‘off-target’, effects of any one mimic.

However, the systematic discovery of synergistic miRNA combinations presents both conceptual and technical challenges. The combinatorial search space grows rapidly with the number of candidate miRNAs, and conventional screening approaches are not optimized to detect nonlinear interactions between regulatory RNAs. Quantitative frameworks for assessing drug synergy have been developed in pharmacology, including models based on additivity, independence, and potency shifts (Yadav et al., 2015; Foucquier and Guedj, 2015; Meyer et al., 2019). Adapting these analytical paradigms to miRNA biology requires integration with high-content phenotypic assays capable of capturing differentiation-related endpoints at scale (Wu et al., 2010; Zhao et al., 2014).

Here, we report the development of a scalable phenotypic screening platform to identify synergistic miRNA combinations that promote neuronal differentiation and growth arrest in neuroblastoma cells. Using SK-N-BE(2)-C neuroblastoma cells and automated imaging-based quantification of neurite outgrowth and confluence, we screened pairwise combinations of candidate differentiation-associated miRNAs at submaximal doses (Zhao et al., 2014; Zhao et al., 2016). Candidate synergistic pairs were identified using the Highest Single Agent framework and subsequently validated through dose–response interaction modeling (Bliss, 1939; Chou, 2006; Meyer et al., 2019).

Through this approach, we identified a robust synergistic interaction between miR-124-3p and miR-363-3p that exceeded expected additive effects and produced substantially greater maximal differentiation phenotypes than either miRNA alone. Mechanistic analysis revealed that these miRNAs regulate complementary target gene sets enriched for pathways involved in neuron projection development, cytoskeletal remodeling, and cell cycle control, supporting a model in which synergy arises from coordinated suppression of independent regulatory bottlenecks (Makeyev et al., 2007; Santos et al., 2016; Agarwal et al., 2015; Subramanian et al., 2005).

## 2. Materials and Methods

### Cell culture

SK-N-BE(2)-C human neuroblastoma cells were obtained from a certified cell repository and maintained in Dulbecco’s Modified Eagle Medium (DMEM) supplemented with 10% fetal bovine serum, 1% penicillin–streptomycin, and L-glutamine at 37 °C in a humidified incubator with 5% CO_2_ (Harenza et al., 2017). Cells were passaged at 70–80% confluence and used for experiments within a limited passage range to ensure phenotypic consistency. Prior to screening, cells were confirmed to be free of mycoplasma contamination.

### miRNA library and reagents

Synthetic miRNA mimics corresponding to candidate differentiation-associated miRNAs were obtained from a commercial supplier and resuspended according to manufacturer specifications. A curated library was assembled based on prior literature implicating these miRNAs in neuronal differentiation, tumor suppression, or neuroblastoma biology (Welch et al., 2007; Chen and Stallings, 2007; Foley et al., 2011; Zhao et al., 2014; Zhao et al., 2016). Negative control miRNA mimics with no known mammalian targets were included on each plate for normalization and quality control.

### Transfection and experimental design

Cells were plated into 384-well plates at a density optimized to allow robust neurite outgrowth and accurate image segmentation over the assay period. Transfections were performed using lipid-mediated delivery reagents according to standardized protocols adapted for high-throughput screening (Du et al., 2013; Wu et al., 2010).

Single-miRNA treatments were performed at a final concentration of 10 nM. Pairwise miRNA combinations were tested at 5 nM + 5 nM to maintain constant total oligonucleotide concentration across all conditions. This design enabled direct comparison between single-agent and combination effects without confounding by total RNA dose. Each condition was tested in technical replicates, and experiments were repeated independently to ensure reproducibility.

### High-content imaging and phenotypic quantification

After transfection, cells were incubated for a defined period to allow induction of differentiation-associated phenotypes. Cells were fixed and stained with nuclear and cytoskeletal markers to enable visualization of cell bodies and neurite projections. Imaging was performed using an Incucyte SX5 live-cell analysis system (Sartorius; software version 2021C) at 10× magnification, with multiple fields captured per well (Wu et al., 2010; Zhao et al., 2014).

Image analysis was conducted using the NeuroTrack analysis module (Sartorius). Cell bodies were segmented based on nuclear and cytoplasmic signals, and neurite projections were identified using skeletonization and tracing algorithms (Wu et al., 2010).

Two primary phenotypic endpoints were extracted:

1. **Neurite outgrowth**, quantified as average neurite length per cell or total neurite length per field.
2. **Cell confluence**, quantified as the fraction of image area occupied by cells, serving as a proxy for growth arrest or cytostatic effects.

These complementary metrics enabled discrimination between differentiation-associated morphological changes and nonspecific cytotoxic responses (Zhao et al., 2014; Zhao et al., 2016). All phenotypic measurements were normalized per plate using a two-point linear transformation. Neurite length was normalized such that the miRNA mimic pool (10 nM) negative control equaled 0 and ATRA (all-trans retinoic acid, 25 µM) equaled 1, using median values at 120 hours: (x − negative control) / (ATRA − negative control). Cell body cluster area was normalized with the miRNA pool as 0 and siPLK1 (10 nM) as 1, using an inverted scale: (siPLK1 − x) / (siPLK1 − miRNA pool).

### Primary synergy analysis using Highest Single Agent (HSA)

For each miRNA pair, the observed phenotypic effect was compared to the maximal effect produced by either miRNA alone at the corresponding concentration using the Highest Single Agent (HSA) framework (Lehár et al., 2009; Foucquier and Guedj, 2015). HSA synergy scores were computed as the difference between the combination response and the stronger single-agent response. Positive deviations from the HSA benchmark were interpreted as evidence of candidate synergistic interactions (Slinker, 1998; Foucquier and Guedj, 2015).

Combinations exceeding predefined HSA thresholds in replicate experiments were classified as candidate synergistic pairs and advanced for secondary validation.

### Dose–response interaction modeling

Candidate synergistic miRNA pairs were subjected to dose–response interaction analysis across a matrix of concentrations for each miRNA. SK-N-BE(2)-C cells were transfected with graded doses of each miRNA individually and in combination, and phenotypic responses were quantified as described above.

Dose–response surfaces were analyzed using zero-interaction potency–based synergy models, including MuSyC-like frameworks, to quantify deviations from predicted non-interacting behavior across the full concentration landscape (Yadav et al., 2015; Meyer et al., 2019; Ivanevski et al., 2022). Metrics extracted from these models included shifts in potency and changes in maximal effect (E_max). Synergy was defined as a statistically significant positive deviation from zero-interaction expectations across replicate experiments.

### Target gene and pathway enrichment analysis

High-confidence target gene lists for miR-124-3p and miR-363-3p were obtained from established miRNA target databases and filtered to include conserved and experimentally supported interactions (Lewis et al., 2005; Agarwal et al., 2015). Overlap between target sets was assessed to determine redundancy and complementarity.

Gene ontology and pathway enrichment analyses were performed using standard bioinformatics tools (Subramanian et al., 2005; Kanehisa et al., 2017; Thomas et al., 2003). Enriched biological processes were identified using corrected significance thresholds, with particular focus on pathways related to neuronal differentiation, cytoskeletal organization, and cell cycle regulation. The GO category “neuron projection development” (GO:0031175) was examined as a representative differentiation-associated pathway regulated by both miRNAs through distinct target subsets.

### Statistical analysis

All experiments were performed with technical and biological replicates. Data were normalized to negative control conditions and reported as mean ± standard error where appropriate. Statistical significance of synergy metrics and phenotypic differences between conditions was assessed using appropriate parametric or nonparametric tests depending on data distribution (Slinker, 1998; Foucquier and Guedj, 2015). Multiple-testing corrections were applied for pathway enrichment analyses (Subramanian et al., 2005). A significance threshold of p < 0.05 was used unless otherwise specified. Statistical analyses were performed using Python 3.9.19 with scipy 1.11.4, pandas 2.0.0, numpy 1.26.4, and matplotlib 3.9.4; dose– response synergy modeling used SynergyFinder 3.0 (Ianevski et al., 2022).

### Statistical analysis of screening data

Pairwise miRNA combinations (n = 946) were screened for effects on neurite length (NL) and cell body cluster area (CBCA) . For neurite length, we calculated an HSA ratio (highest single agent / combination) where values <1.0 indicate synergy. Statistical significance was assessed using Welch’s t-test comparing combination replicates (n = 3) to highest single agent (HSA) replicates (n = 3), using two-sided tests (α = 0.05) to create an unbiased volcano plot showing both synergy and antagonism.

For dual-phenotype filtering, we selected combinations meeting both criteria: (1) neurite length synergy (HSA/combination < 1, p < 0.05 two-sided), and (2) reduced cell body cluster area compared to ATRA (25 µM) (p < 0.05, one-sided test). The ATRA CBCA reference value was computed as the mean of 96 ATRA-treated replicates from the screening data. CBCA was compared using one-sample t-tests against this reference, as we specifically sought combinations with lower clustering than the ATRA control.

Enrichment of CBCA improvement among NL-synergistic combinations was assessed by Fisher’s exact test, comparing the proportion of NL synergies with CBCA < ATRA to the background rate. To formally control FDR, we combined one-sided NL and CBCA p-values per combination using Fisher’s method (chi-squared test with 4 degrees of freedom) and applied Benjamini-Hochberg correction across all 946 combinations. Hits were required to also meet directional criteria (CI < 1 and CBCA mean < ATRA mean).

Label-permutation testing (10,000 iterations) was performed by shuffling CBCA pass/fail labels against fixed NL classifications to estimate the null distribution of dual-positive counts and compute an empirical p-value.

## 3. Results

### Development of a scalable miRNA combination screening platform

To enable systematic discovery of synergistic miRNA interactions, we developed a high-content phenotypic screening platform optimized for quantitative assessment of neuronal differentiation and growth arrest in neuroblastoma cells. The assay was implemented in SK-N-BE(2)-C cells, a well-characterized MYCN-amplified neuroblastoma line that exhibits robust neurite outgrowth in response to differentiation cues and provides a reproducible model for phenotypic screening (Harenza et al., 2017; Zhao et al., 2014; Zhao et al., 2016).

Cells were plated in 384-well format and transfected with individual miRNAs or pairwise combinations using lipid-mediated delivery. Single miRNAs were tested at a final concentration of 10 nM, while combinations were tested at 5 nM + 5 nM to maintain constant total oligonucleotide concentration across conditions. This design allowed direct comparison of single-agent and combination effects while minimizing dose-dependent confounding (Du et al., 2013; Wu et al., 2010).

Differentiation was quantified using automated high-content fluorescence microscopy followed by image-based feature extraction. Two primary phenotypic endpoints were measured: neurite outgrowth, reflecting neuronal differentiation, and cell confluence, reflecting cytostatic or cytotoxic effects. Neurite length was quantified using skeletonization-based algorithms applied to segmented cell bodies and projections, while confluence was computed as the fractional area occupied by cells within each imaging field (**Figure 2**) (Wu et al., 2010; Zhao et al., 2014). These complementary metrics enabled discrimination between true differentiation-associated morphological changes and nonspecific loss of viability (Zhao et al., 2016).

**Figure 2.**
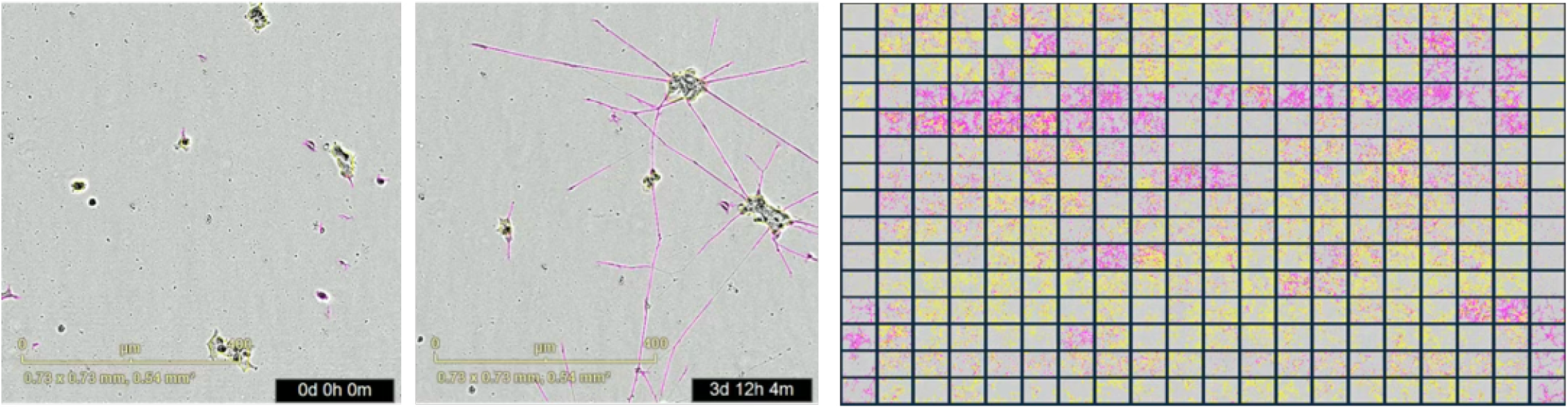
Screening miRNAs for pairs that drive NB cell differentiation. miRNAs implicated in NB differentiation were combinatorially evaluated in SK-N-BE(2)-C cells. Differation was measured with the Incucyte SX5 live cell imaging system and analyzed with Neurotrack. Neurite length and confluence were normalized per plate against positive and negative controls.

A curated library of miRNAs previously implicated in neuronal differentiation and neuroblastoma biology was assembled for combinatorial screening (Welch et al., 2007; Chen and Stallings, 2007; Foley et al., 2011; Zhao et al., 2014; Zhao et al., 2016). All possible pairwise combinations within this library were evaluated in parallel with their corresponding single-miRNA controls. Each condition was tested in technical replicates across independent experimental runs to ensure robustness and reproducibility.

To identify candidate synergistic interactions, primary screening data were analyzed using the Highest Single Agent (HSA) framework. For each miRNA pair, the observed phenotypic effect was compared to the maximal effect achieved by either component miRNA alone at the corresponding concentration. Combinations that exceeded the best-performing single miRNA were classified as exhibiting positive deviation from additivity and advanced for secondary validation. This conservative criterion minimized false-positive synergy calls arising from measurement noise or modest additive effects (Bliss, 1939; Chou, 2006; Slinker, 1998; Foucquier and Guedj, 2015).

Across the screened miRNA pairs, the majority of combinations produced phenotypes consistent with additivity or sub-additivity relative to their component miRNAs. However, a distinct subset of combinations demonstrated marked enhancement of neurite outgrowth beyond the strongest single-agent response. These candidate synergistic pairs were reproducibly observed across replicate experiments and were prioritized for dose– response interaction modeling (Yadav et al., 2015; Meyer et al., 2019).

Together, these results establish a scalable and quantitative platform for phenotypic discovery of synergistic miRNA combinations. The assay integrates high-throughput imaging, conservative synergy triage, and reproducible phenotypic endpoints, providing a foundation for subsequent quantitative validation and mechanistic analysis (Zhao et al., 2014; Zhao et al., 2016).

### Identification of synergistic miRNA pairs

Application of the high-content screening platform to the curated miRNA library yielded a quantitative landscape of phenotypic responses for all tested single miRNAs and their pairwise combinations (**Figure 3**). For each miRNA pair, neurite outgrowth and confluence measurements were normalized to negative control conditions and compared against the corresponding single-miRNA effects using the Highest Single Agent (HSA) model (Bliss, 1939; Chou, 2006; Slinker, 1998; Foucquier and Guedj, 2015).

**Figure 3.**
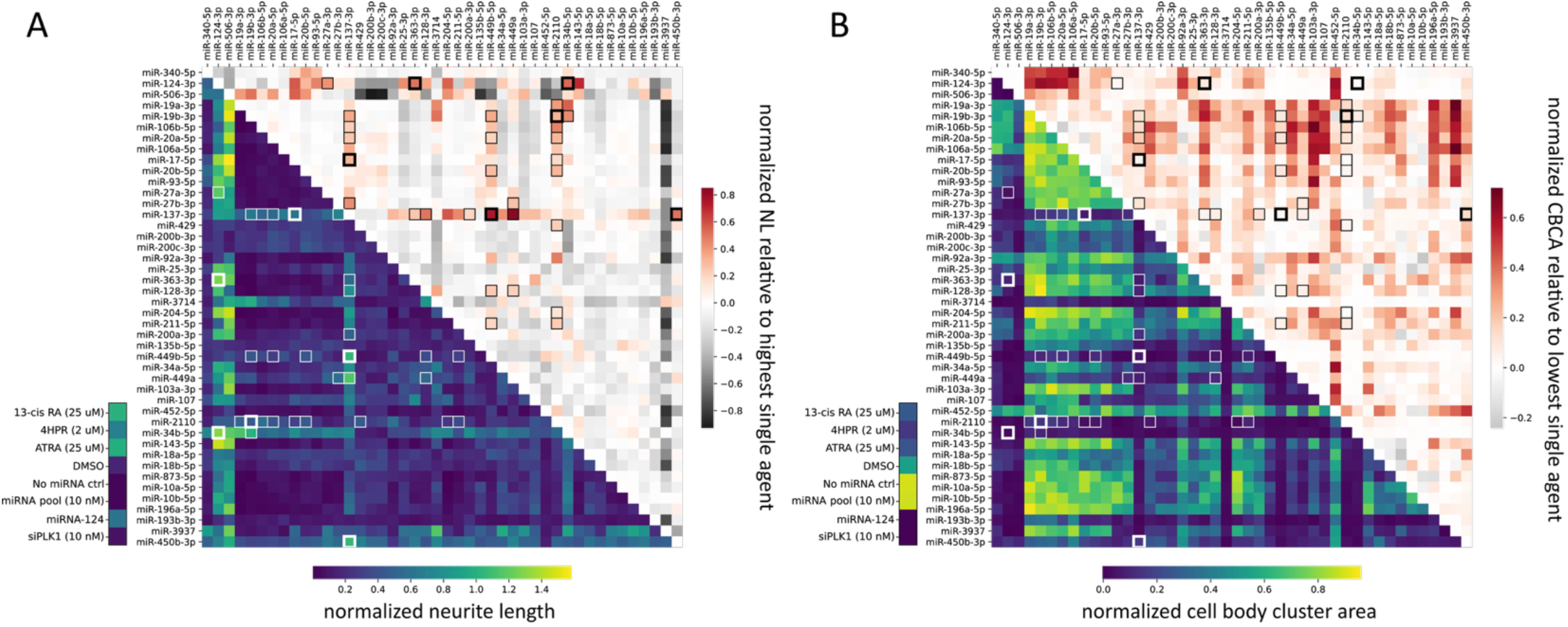
Identifying pairs of miRNAs that drive NB cell differentiation. **(A)** Heatmap of the average of day 4 and day 5 neurite length two-point normalized to ATRA (25 uM) and miRNA pool (10 nM). **(B)** Heatmap of the average of day 4 and day 5 cell body cluster area two-point normalized to siPLK1 and miRNA pool. The lower left-hand triangles of panels A and B depict the normalized metric (neurite length and confluence, left and right, respectively). The upper right-hand triangles depict the difference between the metric and its highest single agent (for neurite length) and lowest single agent (for confluence). miRNAs are clustered by sequence similarity.

The majority of miRNA combinations exhibited phenotypic responses that were comparable to or weaker than the strongest single component miRNA, consistent with additive or sub-additive behavior. This distribution reflects the inherent redundancy and robustness of gene regulatory networks and underscores the challenge of identifying true synergistic interactions within a large combinatorial space (Bartel, 2009; Lewis et al., 2005; Greene et al., 2015). Nevertheless, a discrete subset of miRNA pairs demonstrated reproducible enhancement of neurite outgrowth beyond the maximal effect observed for either miRNA alone, satisfying the HSA criterion for positive synergy.

Candidate synergistic pairs were ranked according to the magnitude of their positive deviation from the HSA benchmark. This prioritization revealed that only a small fraction of tested combinations achieved substantial enhancement of differentiation-associated phenotypes, indicating that synergy is not a generic property of miRNA pairing but instead depends on specific functional complementarity between regulatory RNAs (Al-Lazikani et al., 2012; Agarwal et al., 2015).

Among the highest-ranked interactions, the combination of miR-124-3p and miR-363-3p consistently produced a marked increase in neurite outgrowth relative to either miRNA alone at equivalent total oligonucleotide concentration. This effect was observed across independent experimental replicates and was accompanied by a concomitant reduction in cell confluence, consistent with induction of a differentiated, growth-arrested cellular state rather than nonspecific cytotoxicity (Zhao et al., 2014; Zhao et al., 2016).

Notably, both miR-124-3p and miR-363-3p have previously been implicated individually in neuronal differentiation and tumor suppression. However, when administered as single agents at 10 nM, each produced only partial phenotypic responses, characterized by modest neurite extension and limited growth inhibition. In contrast, their combined administration at 5 nM + 5 nM yielded a phenotype that exceeded the maximal effect of either component miRNA, suggesting cooperative regulation of differentiation-associated pathways (Makeyev et al., 2007; Santos et al., 2016; Nolan et al., 2020; Tivnan et al., 2014).

Additional candidate synergistic pairs were identified within the screening dataset, but the miR-124-3p and miR-363-3p combination exhibited the most robust and reproducible enhancement of the differentiation phenotype and was therefore selected as a representative exemplar for quantitative validation and mechanistic analysis in subsequent experiments (Zhao et al., 2014; Zhao et al., 2016).

These findings demonstrate that while most miRNA combinations behave in an additive or neutral manner, a restricted subset of miRNA pairs can produce emergent phenotypic effects consistent with synergy. The identification of such interactions provides experimental support for the hypothesis that combinatorial miRNA regulation can overcome network-level robustness and induce complex cellular phenotypes more effectively than single-miRNA interventions (Al-Lazikani et al., 2012; Bartel, 2009).

### Dual-phenotype filtering identifies 34 high-confidence synergistic pairs

We tested 946 pairwise miRNA combinations for synergistic effects on neurite outgrowth in SK-N-BE(2) neuroblastoma cells, comparing their activity to ATRA and other retinoid treatments. Because of the large scale of combinatorial screening, we first narrowed the landscape with a single concentration comparison. By comparing combinations of miRNAs (each at 5 nM) against the individual miRNAs at 10 nM, promising candidates can be advanced for higher-resolution dose-response experiments. Comparing combinations with their highest single agent, we identified 49 combinations with significant neurite length synergy (HSA/combination < 1, p < 0.05, two-sided Welch’s t-test; **Figure 5**). Notably, zero of these 49 putative synergistic pairs were from the same miRNA family (vs 3.3% same-family in background), although this depletion did not reach statistical significance (Fisher’s exact p = 0.40) given the low base rate. The direction is nonetheless consistent with the expectation that functionally redundant miRNAs would not synergize.

To identify combinations with multi-phenotype differentiation activity, we applied dual-phenotype filtering requiring combinations to also show significantly reduced cell body clustering compared to ATRA (p < 0.05, one-sided test). Cell body cluster area (CBCA) measures spatial organization of cells, with lower values indicating enhanced differentiation and reduced clustering. NL-synergistic combinations were strongly enriched for CBCA improvement: 82% (40/49) had lower CBCA than the ATRA reference versus 50% background (OR = 4.7, Fisher’s exact p < 0.0001), and 69% (34/49) reached CBCA significance versus a 40% background rate (OR = 3.6, p = 3.3e-5). Label-permutation testing confirmed that the 34 dual-positive hits significantly exceed the null expectation (permutation p < 0.0001, 10,000 iterations).

As a formal multiple-testing correction, we combined NL and CBCA p-values using Fisher’s method and applied Benjamini-Hochberg correction. All 34 dual-positive hits passed at q < 0.05, with 24 passing at q < 0.01, confirming robust combined evidence for both phenotypes. Six dual-positive pairs were subsequently validated by dose-response experiments (6/6 confirmed), providing independent biological confirmation.

### Quantitative validation of synergy by dose–response modeling

To rigorously determine whether the enhanced phenotypic response observed for the miR-124-3p and miR-363-3p combination reflected true synergy rather than a fixed-dose screening artifact, we performed dose–response interaction experiments across a matrix of concentrations for each miRNA. SK-N-BE(2)-C cells were transfected with graded doses of miR-124-3p and miR-363-3p individually and in pairwise combinations, and neurite outgrowth and confluence were quantified using the same high-content imaging pipeline employed in the primary screen (Wu et al., 2010; Zhao et al., 2014; Zhao et al., 2016).

Single-agent dose–response curves revealed that both miR-124-3p and miR-363-3p induced differentiation-associated phenotypes in a concentration-dependent manner but reached limited maximal efficacy when administered alone. Even at higher concentrations, neither miRNA achieved the full extent of neurite outgrowth observed in the initial combination screen, indicating that each miRNA alone was constrained by partial engagement of the underlying differentiation network (Makeyev et al., 2007; Santos et al., 2016; Nolan et al., 2020).

Combination dose–response matrices demonstrated a pronounced upward shift in phenotypic response across a broad range of concentration pairs relative to the expected additive effects derived from single-agent curves. These interaction surfaces were analyzed using zero-interaction potency–based synergy models, which quantify deviations from predicted non-interacting behavior across the full dose–response landscape. This approach allowed discrimination between simple dose additivity and true cooperative interaction (Yadav et al., 2015; Meyer et al., 2019; Ianevski et al., 2022; Foucquier and Guedj, 2015).

Quantitative modeling revealed that the miR-124-3p and miR-363-3p combination exceeded the zero-interaction potency (ZIP) expectation by approximately 20.9%, confirming the presence of a synergistic interaction rather than simple additivity (Bliss, 1939; Chou, 2006; Meyer et al., 2019). Bliss independence analysis using SynergyFinder 3.0 yielded a synergy score of 20.8, with a most synergistic area score of 27.4, independently confirming robust synergy (Ianevski et al., 2022). Moreover, the maximal phenotypic effect (E_max) achieved by the combination was increased by approximately 73% relative to the strongest single-miRNA treatment, demonstrating that synergy manifested not only as a shift in potency but also as a substantial enhancement of maximal differentiation response.

Importantly, the synergistic effect was observed consistently across replicate experiments and was evident in both neurite outgrowth and growth arrest metrics. The combination did not produce disproportionate loss of viability compared with single-miRNA treatments, indicating that the enhanced phenotype reflected coordinated induction of differentiation rather than nonspecific cytotoxicity (Zhao et al., 2014; Zhao et al., 2016).

These results establish that the interaction between miR-124-3p and miR-363-3p represents bona fide synergy across a continuous concentration space. The observed increases in both potency and maximal effect support a model in which the two miRNAs cooperatively regulate distinct but complementary components of the gene regulatory network governing neuronal differentiation and cell cycle exit (Bartel, 2009; Lewis et al., 2005; Agarwal et al., 2015).

Together with the initial HSA-based screen, these dose–response analyses validate the screening platform’s ability to identify true synergistic miRNA combinations and provide a quantitative foundation for subsequent mechanistic interpretation of combinatorial miRNA regulation (Yadav et al., 2015; Meyer et al., 2019).

### Synergistic miRNA combinations induce sustained differentiation and correlate with favorable patient outcomes

To determine whether the synergistic interactions identified in dose–response experiments translated into durable phenotypic changes, we performed time-course analysis of neurite outgrowth and cell body cluster area over an extended 144-hour period following transfection. SK-N-BE(2)-C cells were transfected with representative synergistic miRNA combinations, including miR-124-3p + miR-34b, miR-137 + miR-449b-5p, and miR-137 + miR-450b-3p, and differentiation phenotypes were monitored continuously using live-cell imaging (Wu et al., 2010; Zhao et al., 2014).

All three combinations demonstrated progressive enhancement of neurite outgrowth over the time course, reaching maximal differentiation by approximately 120 hours post-transfection (**Figure 6A**). Notably, the combinations sustained neurite extension throughout the observation period without evidence of phenotypic regression, indicating stable induction of the differentiated state rather than transient morphological changes. In parallel, cell body cluster area measurements revealed progressive reduction in cellular clustering over the same time frame (**Figure 6B**), consistent with coordinated induction of differentiation and growth arrest. The kinetics of these phenotypic changes were consistent across independent synergistic pairs, suggesting a common mechanism of action involving gradual remodeling of the underlying gene regulatory network (Makeyev et al., 2007; Santos et al., 2016).

The sustained differentiation phenotypes induced by synergistic miRNA combinations raised the question of whether endogenous expression of these miRNAs in primary neuroblastoma tumors might correlate with clinical outcomes. To address this, we analyzed publicly available neuroblastoma patient datasets to assess the relationship between miRNA expression levels and overall survival. Patients were stratified into groups based on the number of miRNAs from representative synergistic pairs expressed at high levels (0, 1, or 2 miRNAs) within their tumors.

Kaplan-Meier survival analysis revealed a striking association between coordinated high expression of both miRNAs in a synergistic pair and improved patient outcomes (**Figure 6C**). Patients whose tumors expressed both miRNAs at high levels exhibited significantly better overall survival compared to those with high expression of only one miRNA or neither miRNA (log-rank test, p < 0.05). Importantly, this survival benefit was observed specifically when both miRNAs from a synergistic pair were co-expressed, consistent with the hypothesis that cooperative regulatory activity in patient tumors recapitulates the synergistic phenotypic effects observed in vitro (Welch et al., 2007; Chen and Stallings, 2007; Buechner et al., 2011).

These findings provide three critical insights. First, synergistic miRNA combinations induce durable differentiation phenotypes that persist over biologically relevant time scales, indicating stable reprogramming of cellular state rather than acute pharmacological perturbation. Second, the time-course dynamics suggest that synergistic effects emerge gradually through cumulative suppression of regulatory network components, consistent with the target-space complementarity model. Third, the correlation between endogenous co-expression of synergistic miRNA pairs and favorable clinical outcomes supports the translational relevance of these combinations and suggests that the cooperative regulatory logic identified through phenotypic screening may operate in human neuroblastoma pathogenesis (Matthay et al., 2016; Stallings et al., 2011).

Together, these results establish that synergistic miRNA combinations produce sustained, clinically relevant differentiation phenotypes and provide a mechanistic and translational foundation for subsequent pathway analysis (Zhao et al., 2014; Zhao et al., 2016).

### Target-space and pathway analysis reveals complementary network regulation underlying synergy

To investigate the mechanistic basis of the observed synergy between miR-124-3p and miR-363-3p, we analyzed their predicted and experimentally supported target gene sets and evaluated pathway enrichment associated with each miRNA individually and in combination. This analysis was guided by the hypothesis that synergistic phenotypic effects arise from coordinated regulation of complementary components within the gene regulatory network governing neuronal differentiation and cell cycle control (Bartel, 2009; Lewis et al., 2005; Agarwal et al., 2015).

Target gene lists for miR-124-3p and miR-363-3p were compiled from established miRNA target databases and filtered for high-confidence interactions (TargetScan context++ score < -0.2, conserved sites). miR-124-3p was predicted to target 1,847 genes, while miR-363-3p was predicted to target 1,453 genes. Critically, only 337 targets were shared between the two miRNAs (Jaccard similarity index = 0.18), indicating 82% non-redundancy in their target repertoires. This high degree of target complementarity—where each miRNA regulates largely distinct gene subsets—distinguishes synergistic pairs from redundant combinations and provides a quantitative basis for cooperative regulation. The complementary targets converged on related biological processes despite limited direct overlap, consistent with a model in which synergy emerges from simultaneous suppression of independent regulatory bottlenecks rather than amplification of a single pathway (Greene et al., 2015; Santos et al., 2016).

Gene ontology enrichment analysis demonstrated that targets of both miRNAs were significantly enriched for pathways associated with neuronal differentiation, including neuron projection development, cytoskeletal organization, and regulation of cell morphogenesis (Subramanian et al., 2005; Kanehisa et al., 2017; Thomas et al., 2003). To quantitatively assess whether synergistic pairs achieve complementary pathway regulation, we computed incremental coverage—the additional pathway coverage gained by combining two miRNAs beyond the better single agent. Analysis of 19 synergistic pairs revealed that these combinations exhibited significantly higher incremental coverage of on-target differentiation pathways (mean = 0.082) compared to liability pathways such as apoptosis and ER stress (mean = 0.047, p < 0.001) or housekeeping pathways such as translation and RNA processing (mean = 0.039, p < 0.001) (**Figure 4**). This selective expansion of on-target pathway coverage demonstrates that synergistic miRNA pairs achieve cooperative regulation through complementary targeting of distinct genes within shared biological processes, rather than through redundant suppression of the same targets.

**Figure 4.**
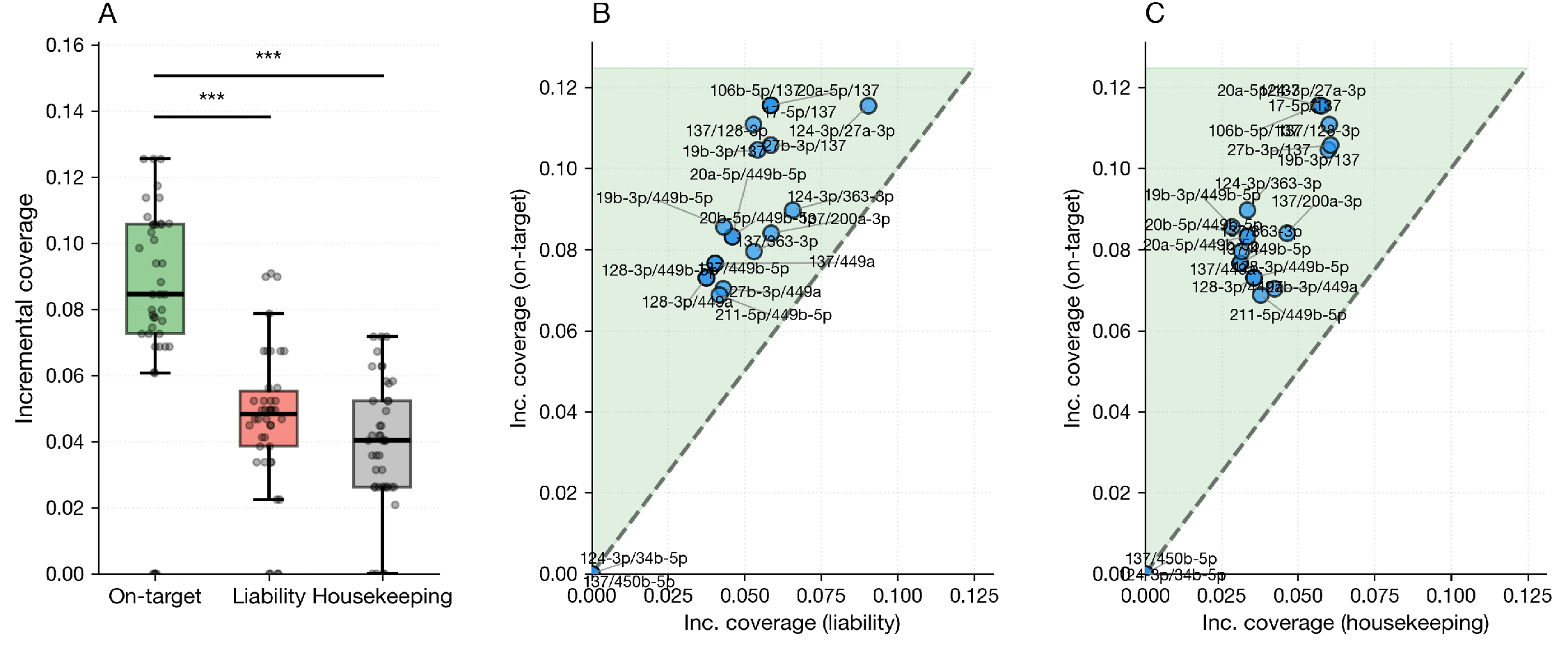
Synergistic miRNA pairs selectively reinforce on-target pathways through complementary target-space coverage. **(A)** Incremental coverage—defined as coverage(A∪B) − max[coverage(A), coverage(B)]—quantifies additional pathway coverage gained by combining two miRNAs with complementary target sets. Higher incremental coverage indicates that the two miRNAs target distinct genes within the same pathway, enabling more complete pathway suppression than either miRNA alone. Synergistic pairs exhibited highest incremental coverage for on-target modules (neurite outgrowth, synapse formation; mean = 0.082), with significantly lower coverage for liability modules (apoptosis, ER stress; mean = 0.047) and housekeeping modules (translation, RNA processing; mean = 0.039) (Mann-Whitney U tests, p < 0.001 for both comparisons, n=19 pairs). Box plots show median, interquartile range, and individual module measurements (points). **(B)** On-target versus liability pathways: All pairs (19/19, 100%) showed greater incremental coverage of on-target versus liability pathways, demonstrating selective expansion of neurite programs through target complementarity while avoiding amplification of toxic pathways. **(C)** On-target versus housekeeping pathways: Similarly, all pairs (19/19, 100%) showed greater incremental coverage of on-target versus housekeeping pathways, demonstrating specificity for neuronal differentiation programs over basic cellular functions. Target predictions from TargetScan v7.2; modules from curated gene sets (GO Biological Process, Reactome, MSigDB Hallmark).

**Figure 5.**
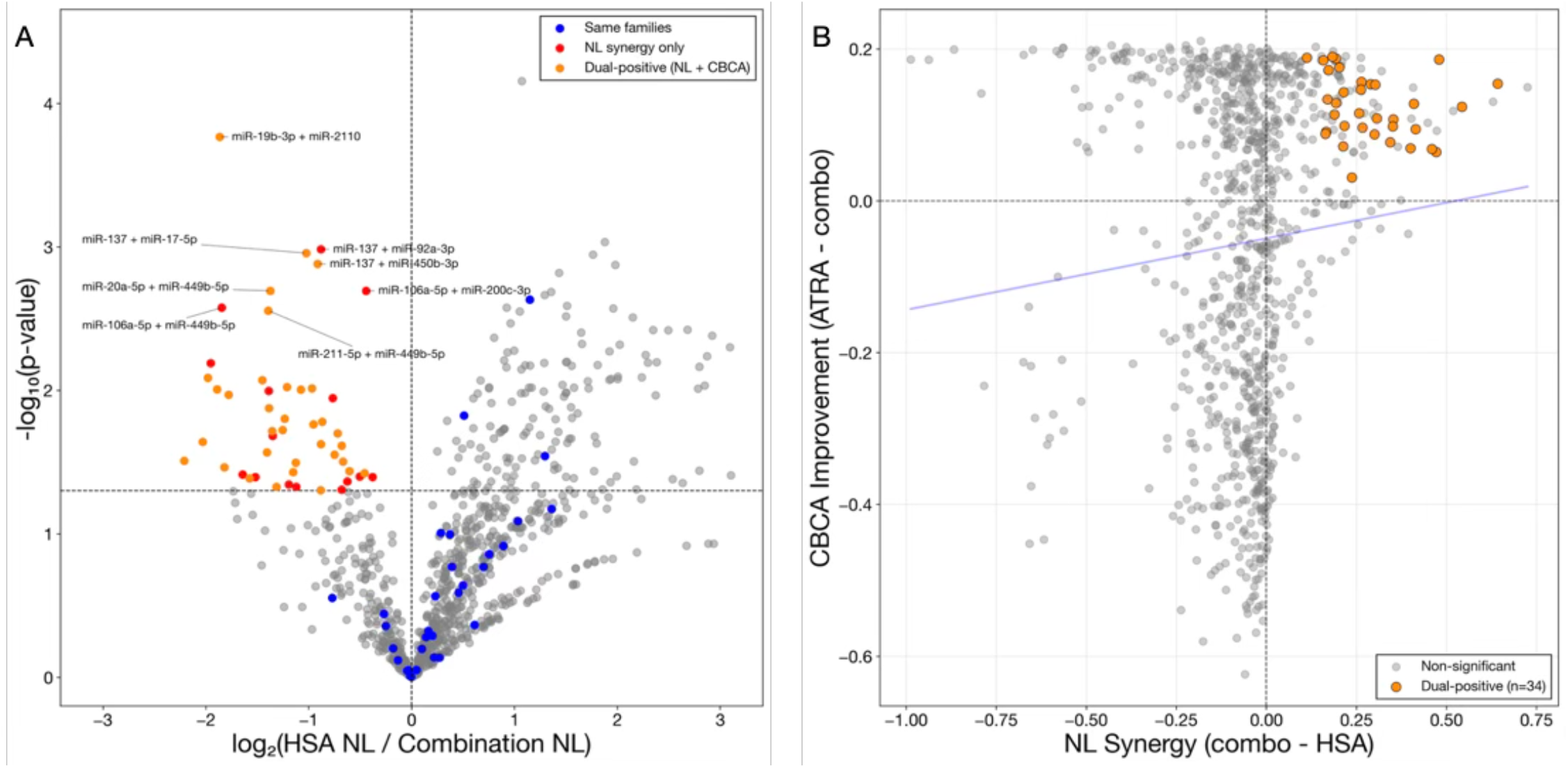
Dual-phenotype filtering identifies neurite length synergies. **(A)** Volcano plot showing 946 pairwise miRNA combinations tested for neurite length synergy versus highest single agent (HSA). X-axis: log_2_(HSA ratio); values < 0 indicate synergy, values > 0 indicate antagonism. Y-axis: -log_10_(p-value) from two-sided Welch’s t-test comparing combination to HSA. Orange points (n=34): dual-positive hits showing both neurite length synergy (CI < 0, p < 0.05) AND significantly reduced cell body cluster area compared to ATRA (p < 0.05, one-sided). Red points: neurite length synergy only. Blue points: 31 same-family miRNA pairs. Gray points: non-significant or antagonistic combinations. Dashed lines indicate significance thresholds (p = 0.05, log_2_(CI) = 0). NL-synergistic pairs are 4.7-fold enriched for directional CBCA improvement (82% vs 50% background, Fisher’s exact p < 0.0001). None of the 49 synergistic pairs (34 dual-positive + 15 NL-only) were from the same miRNA family. All 34 dual-positive hits pass Fisher’s combined test with Benjamini-Hochberg correction at q < 0.05. **(B)** Independence of neurite length and cell body cluster area phenotypes. Scatter plot showing NL synergy (combo - HSA, positive values = synergy) versus CBCA improvement (ATRA - combo, positive values = improved differentiation) for all 946 combinations. Orange points: 34 dual-positive hits from panel A, clustered in the upper-right quadrant (both phenotypes improved). Gray points: non-significant combinations. Linear regression (blue line) demonstrates minimal correlation (Pearson r = 0.081, p = 0.012), R^2^ = 0.007, 99.3% independent variance). NL-synergistic combinations were strongly enriched for directional CBCA improvement (82% vs 50% background, OR = 4.7, Fisher’s exact p < 0.0001), and for statistically significant CBCA improvement (69% vs 40% background, OR = 3.6, p = 3.3e-5). The overlap of 34 dual-positive hits significantly exceeds the permutation null (permutation p < 0.0001, 10,000 iterations).

**Figure 6.**
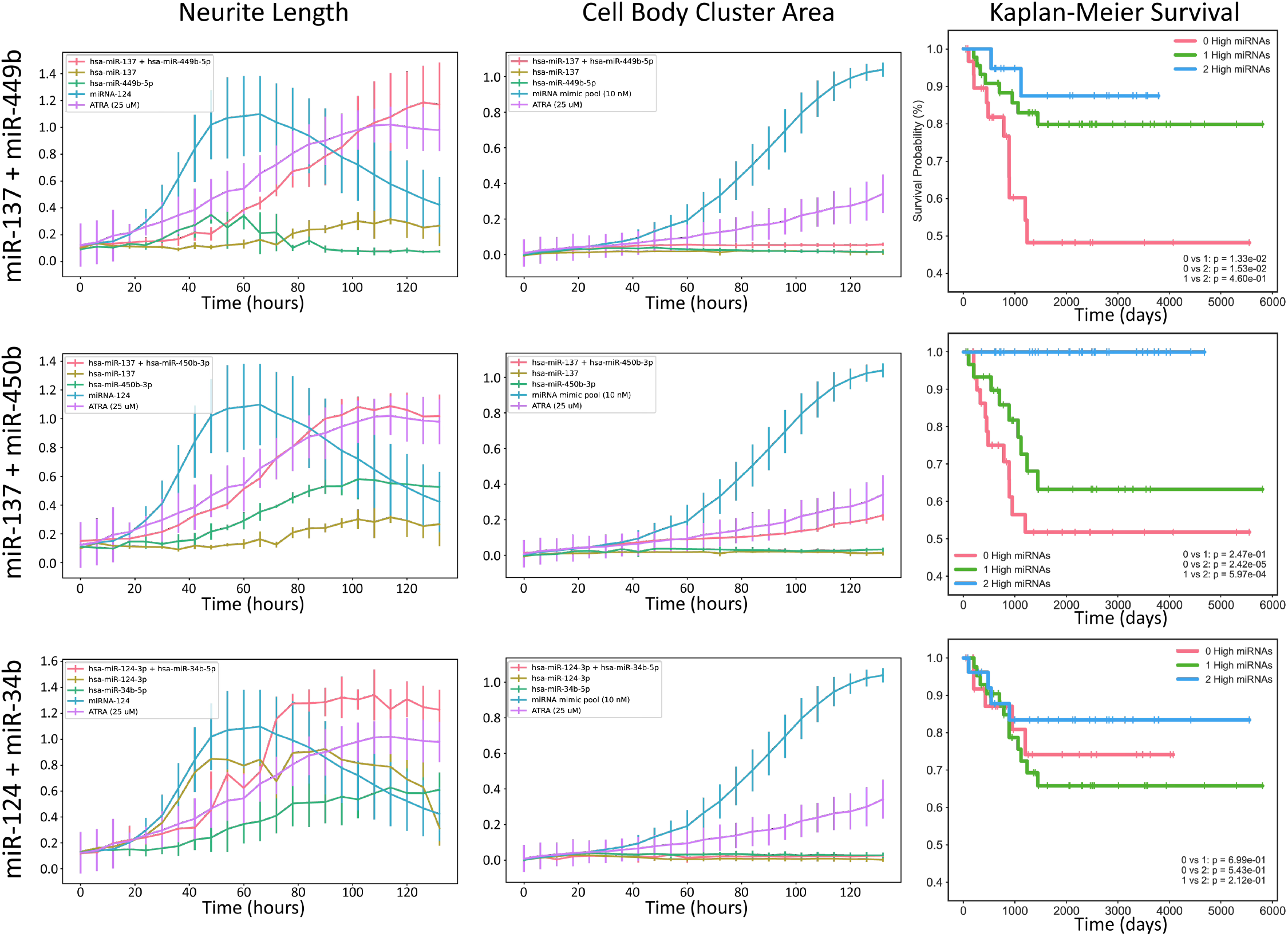
miRNA combinations synergize to drive differentiation and cytostasis of neuroblastoma cells and are strongly associated with positive patient outcomes. **(A)** Neurite length and **(B)** cell body cluster area of NB cells transfected with miRNA mimics (individual and combined, same equimolar amounts) over time. Proliferation monitored by live-cell imaging (IncuCyte) for 144 h (t test; ****, p < 0.0001). Mean±SD, n=3. **(C)** Survival of NB patients grouped by # of miRNAs expressed at high levels (0, 1, or 2).

In addition, miR-124-3p targets showed enrichment for genes involved in neuronal identity and synaptic structure, whereas miR-363-3p targets were preferentially enriched for regulators of cell cycle progression and proliferative signaling. These complementary enrichment profiles suggest that miR-124-3p primarily promotes acquisition of neuronal features, while miR-363-3p concurrently suppresses proliferative programs that antagonize differentiation (Makeyev et al., 2007; Santos et al., 2016; Tivnan et al., 2014; Nolan et al., 2020).

Integration of target-space and phenotypic data revealed that the combination of miR-124-3p and miR-363-3p more effectively reconfigured the transcriptional landscape than either miRNA alone. The combined treatment was associated with enhanced activation of differentiation-associated morphological programs and stronger inhibition of growth-related pathways, consistent with the observed increases in neurite outgrowth and reductions in cell confluence (Zhao et al., 2014; Zhao et al., 2016).

Notably, the enriched gene ontology category of neuron projection development (GO:0031175) encompassed targets from both miRNAs, indicating convergence on a shared biological endpoint through distinct molecular routes. This convergence provides a plausible mechanistic explanation for the observed increase in maximal phenotypic effect under combination treatment: parallel regulation of structural and proliferative determinants of differentiation yields a more complete transition toward a differentiated cellular state (Subramanian et al., 2005; Santos et al., 2016).

These findings support a network-based model of miRNA synergy in which cooperative phenotypic effects arise from complementary regulation of multiple functional modules rather than from intensified repression of a single gene or pathway (Bartel, 2009; Al-Lazikani et al., 2012; Agarwal et al., 2015). By simultaneously engaging differentiation-promoting and proliferation-suppressing programs, the miR-124-3p and miR-363-3p combination achieves a level of phenotypic control that is unattainable by either miRNA individually.

Together, this target-space and pathway analysis provides a mechanistic framework linking quantitative synergy to coordinated network regulation and reinforces the concept of miRNA combinations as programmable modulators of complex cellular phenotypes (Greene et al., 2015; Santos et al., 2016).

## 4. Discussion

This study demonstrates that combinations of miRNAs can act as synergistic regulators of neuronal differentiation in neuroblastoma cells, producing phenotypic effects that exceed those achieved by individual miRNAs alone. By integrating high-content phenotypic screening with quantitative synergy modeling and target-space analysis, we establish a framework for identifying and interpreting cooperative interactions between regulatory RNAs (Zhao et al., 2014; Zhao et al., 2016; Yadav et al., 2015; Meyer et al., 2019). These findings support a paradigm in which miRNA combinations function as programmable network modulators capable of overcoming the robustness and redundancy of oncogenic gene regulatory circuits (Bartel, 2009; Lewis et al., 2005; Greene et al., 2015).

A central insight emerging from this work is that synergy arises not from intensified repression of a single molecular target, but from coordinated regulation of complementary functional modules within the differentiation network. Quantitatively, synergistic miRNA pairs exhibit high target-space complementarity (low Jaccard similarity) coupled with convergent pathway enrichment, enabling more complete suppression of critical biological processes than either miRNA alone. The miR-124-3p and miR-363-3p combination exemplifies this principle: despite sharing only 18% of their predicted targets, both miRNAs converge on neuronal differentiation pathways through distinct molecular routes. miR-124-3p preferentially regulates genes associated with neuronal identity and structural differentiation, whereas miR-363-3p primarily targets regulators of cell cycle progression and proliferative signaling. Together, these miRNAs simultaneously promote acquisition of neuronal features and suppress antagonistic growth programs, yielding a more complete and stable differentiated phenotype (Makeyev et al., 2007; Santos et al., 2016; Nolan et al., 2020; Tivnan et al., 2014). This network-level mechanism of target-space complementarity provides a conceptual explanation for the observed increases in both potency and maximal effect in dose–response synergy analyses (Bliss, 1939; Chou, 2006; Meyer et al., 2019).

The strategy described here parallels the logic of combination therapy in pharmacology, where drugs with distinct but complementary mechanisms are combined to achieve greater efficacy and durability (Al-Lazikani et al., 2012). However, miRNAs offer unique advantages as therapeutic agents in this context. Unlike small molecules or monoclonal antibodies that typically target single proteins, miRNAs intrinsically regulate entire gene programs (He and Hannon, 2004; Bartel, 2009). As a result, combinatorial miRNA therapy enables simultaneous modulation of multiple pathways using a small number of well-chosen regulatory RNAs (Nana-Sinkam and Croce, 2013; van Rooij and Kauppinen, 2014). This property is particularly well suited to differentiation-based therapies, where coordinated shifts in transcriptional state are required rather than isolated pathway inhibition (Reynolds et al., 2003; Matthay et al., 2009; Chlapek et al., 2018).

From a methodological standpoint, this work establishes a scalable approach for discovering synergistic miRNA combinations using conservative triage criteria followed by rigorous quantitative validation. The use of the Highest Single Agent framework as an initial filter minimizes false-positive interactions and allows efficient navigation of the combinatorial search space (Bliss, 1939; Chou, 2006; Slinker, 1998; Foucquier and Guedj, 2015). Subsequent dose–response modeling provides formal confirmation of synergy across concentration ranges, distinguishing true cooperative interactions from artifacts of fixed-dose screening (Yadav et al., 2015; Meyer et al., 2019). This two-tiered strategy can be readily generalized to other cellular phenotypes and disease models, enabling systematic exploration of combinatorial miRNA regulation in diverse biological contexts (Zhao et al., 2014; Zhao et al., 2016).

Despite these advances, several limitations should be acknowledged. First, the current study is confined to in vitro experiments in a single neuroblastoma cell line. While SK-N-BE(2)-C cells provide a robust and well-characterized differentiation model, the generality of the identified synergistic interaction across additional neuroblastoma subtypes and patient-derived models remains to be determined (Harenza et al., 2017; Pinto et al., 2015). Second, the mechanistic analysis relies primarily on target prediction and pathway enrichment rather than direct transcriptomic or proteomic measurements following combination treatment. Future studies integrating RNA sequencing and proteomic profiling will be necessary to map the precise molecular programs engaged by synergistic miRNA pairs (Subramanian et al., 2005; Agarwal et al., 2015). Third, the primary screen used three biological replicates per condition, which provides limited statistical power to detect modest synergistic effects. While this is standard for high-content combinatorial screens of this scale (946 combinations), the resulting false-negative rate may be substantial, and weaker synergistic interactions may have been missed. The six-of-six validation rate in dose-response experiments suggests that the screen’s specificity is high despite the low per-test power.

Another critical challenge lies in therapeutic delivery. Effective and selective delivery of miRNA combinations to tumor cells in vivo remains a major barrier to clinical translation. However, recent advances in lipid nanoparticles, polymeric carriers, and high-density lipoprotein–based delivery systems provide promising avenues for overcoming these obstacles (Babar et al., 2012; Shahzad et al., 2011; Lacko et al., 2015; Raut et al., 2018). The identification of synergistic miRNA pairs with enhanced efficacy may reduce the required dosage of individual miRNAs, potentially improving safety profiles and minimizing off-target effects (van Rooij and Kauppinen, 2014; Nana-Sinkam and Croce, 2013).

The implications of this work extend beyond neuroblastoma. Many cancers and developmental disorders are driven by dysregulation of complex gene networks rather than single dominant mutations (Al-Lazikani et al., 2012; Greene et al., 2015). In such settings, combinatorial miRNA strategies offer a rational means of restoring balanced regulatory control (Bartel, 2009; Santos et al., 2016). More broadly, the framework presented here suggests that miRNA combinations can be designed and optimized as a new class of network therapeutics, analogous to rationally engineered drug combinations but operating at the level of gene regulatory architecture (Nana-Sinkam and Croce, 2013; van Rooij and Kauppinen, 2014).

In summary, our findings provide conceptual and experimental support for miRNA combinations as synergistic agents capable of inducing complex cellular phenotypes through coordinated network regulation. By establishing a reproducible screening and validation platform and demonstrating mechanistic complementarity in a prototypical miRNA pair, this work lays the foundation for the rational development of combinatorial miRNA therapies (Zhao et al., 2014; Zhao et al., 2016; Yadav et al., 2015). Future efforts integrating expanded screening libraries, multi-omics mechanistic analysis, and in vivo validation will be essential to translate this strategy into clinically relevant differentiation-based treatments for neuroblastoma and other diseases characterized by aberrant regulatory networks (Matthay et al., 2016; van Rooij and Kauppinen, 2014).

## Acknowledgments

We acknowledge support from NIH/NINDS (R21NS113344, Penalva), the William and Ella Owens Medical Research Foundation (Penalva, Pertsemlidis), and pilot funds from the Joe R. & Teresa Lozano Long School of Medicine and the Greehey Children’s Cancer Research Institute (Hart, Penalva, Pertsemlidis).

## Declaration of Interests

The authors declare no competing financial interests.

## Data Availability

Study data and analysis code are available upon reasonable request.

## Notes

### Competing Interest Statement

The authors have declared no competing interest.

## References

Althoff, K., A. Beckers, A. Odersky, P. Mestdagh, J. Koster, I. M. Bray, K. Bryan, J. Vandesompele, F. Speleman, R. L. Stallings, A. Schramm, A. Eggert, A. Sprussel and J. H. Schulte (2013). “MiR-137 functions as a tumor suppressor in neuroblastoma by downregulating KDM1A.” Int J Cancer 133(5): 1064–1073.

Behlke, M. A. (2008). “Chemical modification of siRNAs for in vivo use.” Oligonucleotides 18(4): 305–319.

Bray, I., A. Tivnan, K. Bryan, N. H. Foley, K. M. Watters, L. Tracey, A. M. Davidoff and R. L. Stallings (2011). “MicroRNA-542-5p as a novel tumor suppressor in neuroblastoma.” Cancer Lett 303(1): 56–64.

Chen, Y. and R. L. Stallings (2007). “Differential patterns of microRNA expression in neuroblastoma are correlated with prognosis, differentiation, and apoptosis.” Cancer Res 67(3): 976–983.

Chlapek, P., V. Slavikova, P. Mazanek, J. Sterba and R. Veselska (2018). “Why differentiation therapy sometimes fails: Molecular mechanisms of resistance to retinoids.” Int J Mol Sci 19(1).

Cohn, S. L., A. D. Pearson, W. B. London, T. Monclair, P. F. Ambros, G. M. Brodeur, A. Faldum, B. Hero, T. Iehara, D. Machin, V. Mosseri, T. Simon, A. Garaventa, V. Castel, K. K. Matthay and I. T. Force (2009). “The International Neuroblastoma Risk Group (INRG) classification system: an INRG Task Force report.” J Clin Oncol 27(2): 289–297.

Corey, D. R. (2007). “Chemical modification: the key to clinical application of RNA interference?” J Clin Invest 117(12): 3615–3622.

Crooke, S. T., J. L. Witztum, C. F. Bennett and B. F. Baker (2018). “RNA-targeted therapeutics.” Cell Metab 27(4): 714–739.

Elmen, J., M. Lindow, S. Schutz, M. Lawrence, A. Petri, S. Obad, M. Lindholm, M. Hedtjarn, H. F. Hansen, U. Berger, S. Gullans, P. Kearney, P. Sarnow, E. M. Straarup and S. Kauppinen (2008). “LNA-mediated microRNA silencing in non-human primates.” Nature 452(7189): 896–899.

Evangelisti, C., M. C. Florian, I. Massimi, C. Dominici, G. Giannini, S. Galardi, M. C. Bue, S. Massalini, H. P. McDowell, E. Messi, A. Gulino, M. G. Farace and S. A. Ciafre (2009). “MiR-128 up-regulation inhibits Reelin and DCX expression and reduces neuroblastoma cell motility and invasiveness.” FASEB J 23(12): 4276–4287.

Foucquier, J. and M. Guedj (2015). “Analysis of drug combinations: current methodological landscape.” Pharmacol Res Perspect 3(3): e00149.

Greene, C. S., A. Krishnan, A. K. Wong, E. Ricciotti, R. A. Zelaya, D. S. Himmelstein, R. Zhang, B. M. Hartmann, E. Zaslavsky, S. C. Sealfon, D. I. Chasman, G. A. FitzGerald, K. Dolinski, T. Grosser and O. G. Troyanskaya (2015). “Understanding multicellular function and disease with human tissue-specific networks.” Nat Genet 47(6): 569–576.

Han, D., X. Dong, D. Zheng and J. Nao (2019). “MiR-124 and the underlying therapeutic promise of neurodegenerative disorders.” Front Pharmacol 10: 1555.

Hu, B., L. Zhong, Y. Weng, L. Peng, Y. Huang, Y. Zhao and X. J. Liang (2020). “Therapeutic siRNA: state of the art.” Signal Transduct Target Ther 5(1): 101.

Hwang, J., C. Chang, J. H. Kim, C. T. Oh, H. N. Lee, C. Lee, D. Oh, C. Lee, B. Kim, S. W. Hong and D. K. Lee (2016). “Development of cell-penetrating asymmetric interfering RNA targeting connective tissue growth Factor.” J Invest Dermatol 136(11): 2305–2313.

Ianevski, A., A. K. Giri and T. Aittokallio (2022). “SynergyFinder 3.0: an interactive analysis and consensus interpretation of multi-drug synergy at the population and sample levels.” Nucleic Acids Res 50(W1): W587–W595.

Kosti, A., R. Barreiro, G. D. A. Guardia, S. Ostadrahimi, E. Kokovay, A. Pertsemlidis, P. A. F. Galante and L. O. F. Penalva (2021). “Synergism of proneurogenic miRNAs provides a more effective strategy to target glioma stem cells.” Cancers (Basel) 13(2).

Kosti, A., L. Du, H. Shivram, M. Qiao, S. Burns, J. G. Garcia, A. Pertsemlidis, V. R. Iyer, E. Kokovay and L. O. F. Penalva (2020). “ELF4 Is a target of miR-124 and promotes neuroblastoma proliferation and undifferentiated state.” Mol Cancer Res 18(1): 68–78.

Lehár, J., A. Krueger, W. Avery, A. M. Heilbut, L. M. Johansen, E. R. Price, R. J. Rickles, G. F. Short III, J. E. Staunton, X. Jin, M. S. Lee, G. R. Zimmermann and A. A. Borisy (2009). “Synergistic drug combinations tend to improve therapeutically relevant selectivity.” Nat Chem Biol 5(9): 674–681.

Lo-Coco, F., G. Avvisati, M. Vignetti, C. Thiede, S. M. Orlando, S. Iacobelli, F. Ferrara, P. Fazi, L. Cicconi, E. Di Bona, G. Specchia, S. Sica, M. Divona, A. Levis, W. Fiedler, E. Cerqui, M. Breccia, G. Fioritoni, H. R. Salih, M. Cazzola, L. Melillo, A. M. Carella, C. H. Brandts, E. Morra, M. von Lilienfeld-Toal, B. Hertenstein, M. Wattad, M. Lubbert, M. Hanel, N. Schmitz, H. Link, M. G. Kropp, A. Rambaldi, G. La Nasa, M. Luppi, F. Ciceri, O. Finizio, A. Venditti, F. Fabbiano, K. Dohner, M. Sauer, A. Ganser, S. Amadori, F. Mandelli, H. Dohner, G. Ehninger, R. F. Schlenk, U. Platzbecker, d. A. Gruppo Italiano Malattie Ematologiche, G. German-Austrian Acute Myeloid Leukemia Study and L. Study Alliance (2013). “Retinoic acid and arsenic trioxide for acute promyelocytic leukemia.” N Engl J Med 369(2): 111–121.

Mahmoudi, E. and M. J. Cairns (2017). “MiR-137: an important player in neural development and neoplastic transformation.” Mol Psychiatry 22(1): 44–55.

Makeyev, E. V., J. Zhang, M. A. Carrasco and T. Maniatis (2007). “The microRNA miR-124 promotes neuronal differentiation by triggering brain-specific alternative pre-mRNA splicing.” Mol Cell 27(3): 435–448.

Manoharan, M. (2004). “RNA interference and chemically modified small interfering RNAs.” Curr Opin Chem Biol 8(6): 570–579.

Matthay, K. K., J. G. Villablanca, R. C. Seeger, D. O. Stram, R. E. Harris, N. K. Ramsay, P. Swift, H. Shimada, C. T. Black, G. M. Brodeur, R. B. Gerbing and C. P. Reynolds (1999). “Treatment of high-risk neuroblastoma with intensive chemotherapy, radiotherapy, autologous bone marrow transplantation, and 13-cis-retinoic acid. Children’s Cancer Group.” N Engl J Med 341(16): 1165–1173.

Nolan, J. C., M. Salvucci, S. Carberry, A. Barat, M. F. Segura, J. Fenn, J. H. M. Prehn, R. L. Stallings and O. Piskareva (2020). “A context-dependent role for miR-124-3p on cell phenotype, viability and chemosensitivity in neuroblastoma in vitro.” Front Cell Dev Biol 8: 559553.

Osborn, M. F. and A. Khvorova (2018). “Improving siRNA delivery in vivo through lipid conjugation.” Nucleic Acid Ther 28(3): 128–136.

Park, J. R., R. Bagatell, S. L. Cohn, A. D. Pearson, J. G. Villablanca, F. Berthold, S. Burchill, A. Boubaker, K. McHugh, J. G. Nuchtern, W. B. London, N. L. Seibel, O. W. Lindwasser, J. M. Maris, P. Brock, G. Schleiermacher, R. Ladenstein, K. K. Matthay and D. Valteau-Couanet (2017). “Revisions to the International Neuroblastoma Response Criteria: A Consensus Statement From the National Cancer Institute Clinical Trials Planning Meeting.” J Clin Oncol 35(22): 2580–2587.

Pinto, N. R., M. A. Applebaum, S. L. Volchenboum, K. K. Matthay, W. B. London, P. F. Ambros, A. Nakagawara, F. Berthold, G. Schleiermacher, J. R. Park, D. Valteau-Couanet, A. D. Pearson and S. L. Cohn (2015). “Advances in risk classification and treatment strategies for neuroblastoma.” J Clin Oncol 33(27): 3008–3017.

Prakash, T. P., A. M. Kawasaki, E. V. Wancewicz, L. Shen, B. P. Monia, B. S. Ross, B. Bhat and M. Manoharan (2008). “Comparing in vitro and in vivo activity of 2’-O-[2-(methylamino)-2-oxoethyl]-and 2’-O-methoxyethyl-modified antisense oligonucleotides.” J Med Chem 51(9): 2766–2776.

Santos, M. C., A. N. Tegge, B. R. Correa, S. Mahesula, L. Q. Kohnke, M. Qiao, M. A. Ferreira, E. Kokovay and L. O. Penalva (2016). “miR-124, -128, and -137 orchestrate neural differentiation by acting on overlapping gene sets containing a highly connected transcription tactor network.” Stem Cells 34(1): 220–232.

Simoes-Costa, M. and M. E. Bronner (2015). “Establishing neural crest identity: a gene regulatory recipe.” Development 142(2): 242–257.

Slinker, B. K. (1998). “The statistics of synergism.” J Mol Cell Cardiol 30(4): 723–731.

Springer, A. D. and S. F. Dowdy (2018). “GalNAc-siRNA conjugates: Leading the way for delivery of RNAi therapeutics.” Nucleic Acid Ther 28(3): 109–118.

Stallings, R. L. (2009). “MicroRNA involvement in the pathogenesis of neuroblastoma: potential for microRNA mediated therapeutics.” Curr Pharm Des 15(4): 456–462.

Stallings, R. L., N. H. Foley, I. M. Bray, S. Das and P. G. Buckley (2011). “MicroRNA and DNA methylation alterations mediating retinoic acid induced neuroblastoma cell differentiation.” Semin Cancer Biol 21(4): 283–290.

Sun, G., P. Ye, K. Murai, M. F. Lang, S. Li, H. Zhang, W. Li, C. Fu, J. Yin, A. Wang, X. Ma and Y. Shi (2011). “miR-137 forms a regulatory loop with nuclear receptor TLX and LSD1 in neural stem cells.” Nat Commun 2: 529.

Tamim, S., D. T. Vo, P. J. Uren, M. Qiao, E. Bindewald, W. K. Kasprzak, B. A. Shapiro, H. I. Nakaya, S. C. Burns, P. R. Araujo, I. Nakano, A. J. Radek, S. Kuersten, A. D. Smith and L. O. Penalva (2014). “Genomic analyses reveal broad impact of miR-137 on genes associated with malignant transformation and neuronal differentiation in glioblastoma cells.” PLoS One 9(1): e85591.

Tan, C. L., J. L. Plotkin, M. T. Veno, M. von Schimmelmann, P. Feinberg, S. Mann, A. Handler, J. Kjems, D. J. Surmeier, D. O’Carroll, P. Greengard and A. Schaefer (2013). “MicroRNA-128 governs neuronal excitability and motor behavior in mice.” Science 342(6163): 1254–1258.

Thatcher, J. E. and N. Isoherranen (2009). “The role of CYP26 enzymes in retinoic acid clearance.” Expert Opin Drug Metab Toxicol 5(8): 875–886.

van Groningen, T., J. Koster, L. J. Valentijn, D. A. Zwijnenburg, N. Akogul, N. E. Hasselt, M. Broekmans, F. Haneveld, N. E. Nowakowska, J. Bras, C. J. M. van Noesel, A. Jongejan, A. H. van Kampen, L. Koster, F. Baas, L. van Dijk-Kerkhoven, M. Huizer-Smit, M. C. Lecca, A. Chan, A. Lakeman, P. Molenaar, R. Volckmann, E. M. Westerhout, M. Hamdi, P. G. van Sluis, M. E. Ebus, J. J. Molenaar, G. A. Tytgat, B. A. Westerman, J. van Nes and R. Versteeg (2017). “Neuroblastoma is composed of two super-enhancer-associated differentiation states.” Nat Genet 49(8): 1261–1266.

Veal, G. J., J. Errington, S. E. Rowbotham, N. A. Illingworth, G. Malik, M. Cole, A. K. Daly, A. D. Pearson and A. V. Boddy (2013). “Adaptive dosing approaches to the individualization of 13-cis-retinoic acid (isotretinoin) treatment for children with high-risk neuroblastoma.” Clin Cancer Res 19(2): 469–479.

Wahlestedt, C., P. Salmi, L. Good, J. Kela, T. Johnsson, T. Hokfelt, C. Broberger, F. Porreca, J. Lai, K. Ren, M. Ossipov, A. Koshkin, N. Jakobsen, J. Skouv, H. Oerum, M. H. Jacobsen and J. Wengel (2000). “Potent and nontoxic antisense oligonucleotides containing locked nucleic acids.” Proc Natl Acad Sci U S A 97(10): 5633–5638.

Ward, E., C. DeSantis, A. Robbins, B. Kohler and A. Jemal (2014). “Childhood and adolescent cancer statistics, 2014.” CA Cancer J Clin 64(2): 83–103.

Yoo, B. H., E. Bochkareva, A. Bochkarev, T. C. Mou and D. M. Gray (2004). “2’-O-methyl-modified phosphorothioate antisense oligonucleotides have reduced non-specific effects in vitro.” Nucleic Acids Res 32(6): 2008–2016.

Yu, A. L., A. L. Gilman, M. F. Ozkaynak, W. B. London, S. G. Kreissman, H. X. Chen, M. Smith, B. Anderson, J. G. Villablanca, K. K. Matthay, H. Shimada, S. A. Grupp, R. Seeger, C. P. Reynolds, A. Buxton, R. A. Reisfeld, S. D. Gillies, S. L. Cohn, J. M. Maris, P. M. Sondel and G. Children’s Oncology (2010). “Anti-GD2 antibody with GM-CSF, interleukin-2, and isotretinoin for neuroblastoma.” N Engl J Med 363(14): 1324–1334.

Zhao, Z., X. Ma, T. H. Hsiao, G. Lin, A. Kosti, X. Yu, U. Suresh, Y. Chen, G. E. Tomlinson, A. Pertsemlidis and L. Du (2014). “A high-content morphological screen identifies novel microRNAs that regulate neuroblastoma cell differentiation.” Oncotarget 5(9): 2499–2512.

Zhao, Z., X. Ma, S. D. Shelton, D. C. Sung, M. Li, D. Hernandez, M. Zhang, M. D. Losiewicz, Y. Chen, A. Pertsemlidis, X. Yu, Y. Liu and L. Du (2016). “A combined gene expression and functional study reveals the crosstalk between N-Myc and differentiation-inducing microRNAs in neuroblastoma cells.” Oncotarget 7(48): 79372–79387.

Zhao, Z., X. Ma, D. Sung, M. Li, A. Kosti, G. Lin, Y. Chen, A. Pertsemlidis, T. H. Hsiao and L. Du (2015). “microRNA-449a functions as a tumor suppressor in neuroblastoma through inducing cell differentiation and cell cycle arrest.” RNA Biol 12(5): 538–554.

Zhao, Z., V. Partridge, M. Sousares, S. D. Shelton, C. L. Holland, A. Pertsemlidis and L. Du (2018). “microRNA-2110 functions as an onco-suppressor in neuroblastoma by directly targeting Tsukushi.” PLoS One 13(12): e0208777.

